# The mitotic spindle is chiral due to torques generated by motor proteins

**DOI:** 10.1101/167437

**Authors:** Maja Novak, Bruno Polak, Juraj Simunić, Zvonimir Boban, Barbara Kuzmić, Andreas W. Thomae, Iva M. Tolić, Nenad Pavin

## Abstract

Mitosis relies on forces generated in the spindle, a micro-machine composed of microtubules and associated proteins^1,2^. Forces are required for the congression of chromosomes to the metaphase plate and separation of chromatids in anaphase^3-6^. However, torques may also exist in the spindle, yet they have not been investigated. Here we show that the spindle is chiral. Chirality is evident from the finding that microtubule bundles follow a left-handed helical path, which cannot be explained by forces but rather by torques acting in the bundles. STED super-resolution microscopy, as well as confocal microscopy, of human spindles shows that the bundles have complex curved shapes. The average helicity of the bundles with respect to the spindle axis is 1.2°/μm. Inactivation of kinesin-5 (Eg5/Kif11) abolished the chirality of the spindle, suggesting that this motor generates the helical shape of microtubule bundles. To explain the observed shapes, we introduce a theoretical model for the balance of forces and torques acting in the spindle, and show that torque is required to generate the helical shapes. We conclude that torques generated by motor proteins, in addition to forces, exist in the spindle and determine its architecture.

---

We set out to infer torques and forces in the spindle, by using the shape of microtubule bundles. We first used stimulated emission depletion (STED) super-resolution microscopy^7,8^ to determine the shapes of microtubule bundles in metaphase spindles. We used HeLa cells with labeled microtubules, kinetochores and centrioles (Fig. 1a) and U2OS cells with labeled microtubules and kinetochores (Extended Data Fig. 1a). Single optical sections of metaphase spindles showed that microtubule bundles are continuous almost from pole to pole and acquire complex curved shapes (Fig. 1a). Typically, the outer bundles are curved and resemble a C-letter shape. Interestingly, some bundles throughout the spindle resemble the letter S. Thus, STED images of the spindle suggest that microtubules are arranged into well-defined bundles exhibiting a variety of shapes, which run continuously almost through the whole spindle.

**Figure 1.**
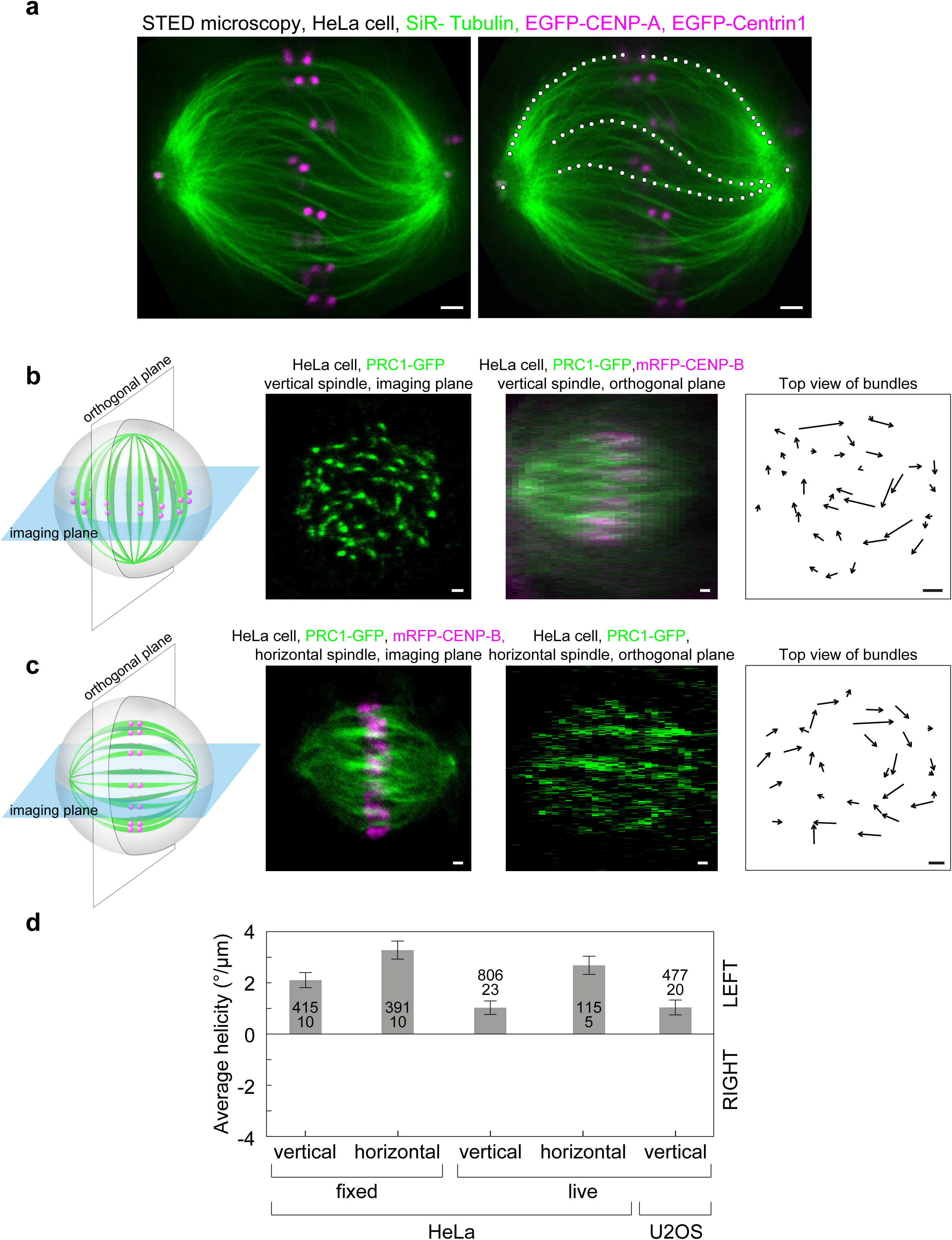
Mitotic spindle has chiral structure. **a,** STED image of the metaphase spindle in HeLa cell expressing EGFP-CENP-A and EGFP-centrin1 (magenta), microtubules labelled with SiR-tubulin (green) (left). Different shapes of microtubule bundles (white tracks) extend almost through the whole spindle (left). **b,** From left to right: scheme of imaging of a vertically oriented spindle; equatorial plane of the vertical spindle in a HeLa cell expressing PRC1-GFP and chromosomes labelled with SiR-DNA (Only PRC1-GFP signal is shown for clarity); projection of the same spindle onto a horizontal plane (PRC1-green, DNA-magenta); arrows connecting starting and ending points of PRC1-GFP bundles traced in the upwards direction. Note that the bundles rotate clockwise, revealing left-handed chirality of the spindle. **c,** From left to right: scheme of imaging of a horizontally oriented spindle; equatorial plane of the horizontal spindle (legend as in **b**); projection of the same spindle onto a vertical plane; arrows connecting starting and ending points of PRC1-GFP bundles traced in the upwards direction. **d,** Bars show spindle helicity, calculated over all bundles for different conditions and cell lines, as the average change in angle with height. Positive values denote left-handed helicity, negative values denote right-handed helicity. Data are shown for HeLa cells expressing PRC1-GFP, and U2OS cells expressing CENP-A-GFP, mCherry-α-tubulin and photoactivatable (PA)-GFP-tubulin. The numbers in each bar represent the number of bundles (top) and cells (bottom), error bars represent s.e.m. Scale bars in panels **a-c** represent 1μm.

In order to obtain a complete three-dimensional contour of microtubule bundles, we used vertically oriented spindles, which are found occasionally in a population of mitotic cells, and imaged them by confocal microscopy (Fig. 1b). In these spindles, optical sections are roughly perpendicular to the microtubule bundles, allowing for precise determination of the bundle location in each section and thus of the whole three-dimensional contour (Methods). We used fixed HeLa cells expressing GFP-tagged protein regulator of cytokinesis 1 (PRC1)^9,10^, which binds overlap zones of antiparallel microtubules^11,12^. Almost all overlap bundles in the spindle are associated with a pair of sister kinetochores, acting as a bridge between the respective kinetochore fibers^13-15^. Thus, PRC1-GFP shows the position of overlap bundles together with their kinetochore fibres, without interference from other microtubules such as polar and astral ones. PRC1-labeled bundles, which appear as spots in a single image plane, were tracked through the z-stacks (Fig. 1b and Methods). We found that individual bundles follow a left-handed helical path along the spindle axis (Fig. 1b and Supplementary Video 1). The arrows connecting starting and ending points of PRC1-GFP bundles traced in the upwards direction rotate clockwise, revealing left-handed chirality of the spindle (Fig. 1b and Extended Data Fig 1b). A 3D plot of all traced points shows the helical shape and spatial arrangement of the bundles (Supplementary Video 2). The helicity of bundles, defined as the average change in angle with height where positive numbers denote left-handed helicity, was 2.1±0.3 °/μm (results are mean±s.e.m. averaged over bundles, unless stated otherwise; n=415 bundles from 10 cells; Fig. 1d). Similar results were obtained when averaging the helicity over cells (Extended Data Fig. 1d). We conclude that the mitotic spindle is a chiral object with left-handed helicity of the microtubule bundles.

To test whether chirality is a specific property of vertically oriented spindles, we measured the average helicity of bundles in fixed horizontally oriented spindles (Fig. 1c). To trace the bundles in these spindles, we rearranged the z-stacks to obtain the slices perpendicular to the spindle axis, similar to the z-stacks of vertical spindles (Fig. 1c, Extended Data Fig 1c and Methods). The bundles in horizontally oriented spindles showed left-handed helicity as in vertical spindles, but with a larger value of 3.3±0.4 °/μm (n=391 bundles from 10 cells; Fig. 1d, Extended Data Fig. 1d shows results averaged over cells). Thus, spindles have left-handed chirality independent of its orientation within the cell.

To test if the fixation process causes spindle chirality, we imaged both vertical and horizontal spindles in live HeLa cells expressing PRC1-GFP. In both cases we observed the left-handed chiral nature of the spindle with helicities for vertical and horizontal spindles of 1.0±0.3 °/μm (n=806 bundles from 23 cells) and 2.7±0.4 °/μm (115 bundles from 5 cells), respectively (Fig. 1d, Extended Data Fig. 1d shows results averaged over cells). These results indicate that chirality is a property of spindles in live cells as well as in fixed cells. Finally, to test if the chirality is limited to a specific cell line, we examined the spindles in U2OS cells expressing mCherry-tubulin and GFP-CENP-A. The helicity of the bundles in these spindles was 1.0±0.3 °/μm (n=477 bundles from 20 cells; Fig. 1d, Extended Data Fig. 1d shows results averaged over cells) indicating that in human cells, microtubule bundles follow a left-handed helical path in different cell lines. Taken together, our results suggest that even though the average helicity value differs among various conditions and cell lines, the bundles consistently twist in a left-handed direction with average value for all cells of 1.2±0.3 °/μm. We conclude that left-handed chirality is a robust feature of the spindle.

Next, we set out to investigate the molecular mechanisms that contribute to the generation of twist in the spindle. We first hypothesized that twist is generated within the bundles, by motor proteins that rotate the microtubule while walking. In vitro gliding motility studies showed that kinesin-5 (Kif11/Eg5) switches protofilaments with biased steps towards the left^16^. To inactivate Eg5, we treated HeLa cells with vertically oriented spindles in metaphase with S-trityl-L-cysteine (STLC), a reversible tight-binding inhibitor^17,18^. The arrows connecting starting and ending points of the bundles traced in the upwards direction show a change from a clockwise rotation before treatment to a more random distribution after STLC treatment (Fig. 2a). Inactivation of Eg5 caused the reduction of bundle helicity in the same cells from 0.6±0.3 °/μm (n=347 bundles from 10 cells) before STLC treatment to 0.2±0.3 °/μm (n=341 bundles from 10 cells) and 0.2±0.3 °/μm (n=153 bundles from 6 cells) at 5 and 10 minutes after STLC addition, respectively (Fig. 2b, 2c, Extended Data Fig. 2a shows results averaged over cells). Whereas the helicities before STLC treatment were significantly different from 0 (p=4x10^−7^), no significant difference from 0 was found after STLC addition (p=0.11 and 0.28 for 5 and 10 minutes after STLC addition, respectively). Thus, the average bundle twist was left-handed before the treatment, whereas there was no preferred handedness after Eg5 inactivation. On the contrary, the bundle helicity in control cells did not change significantly within 10 minutes, suggesting that the helicity does not fluctuate significantly during metaphase in individual cells (Fig. 2d). Similarly, Eg5 inactivation in U2OS abolished spindle chirality (Fig.2c). Based on these results, we conclude that kinesin-5 contributes to the generation of spindle chirality.

**Figure 2.**
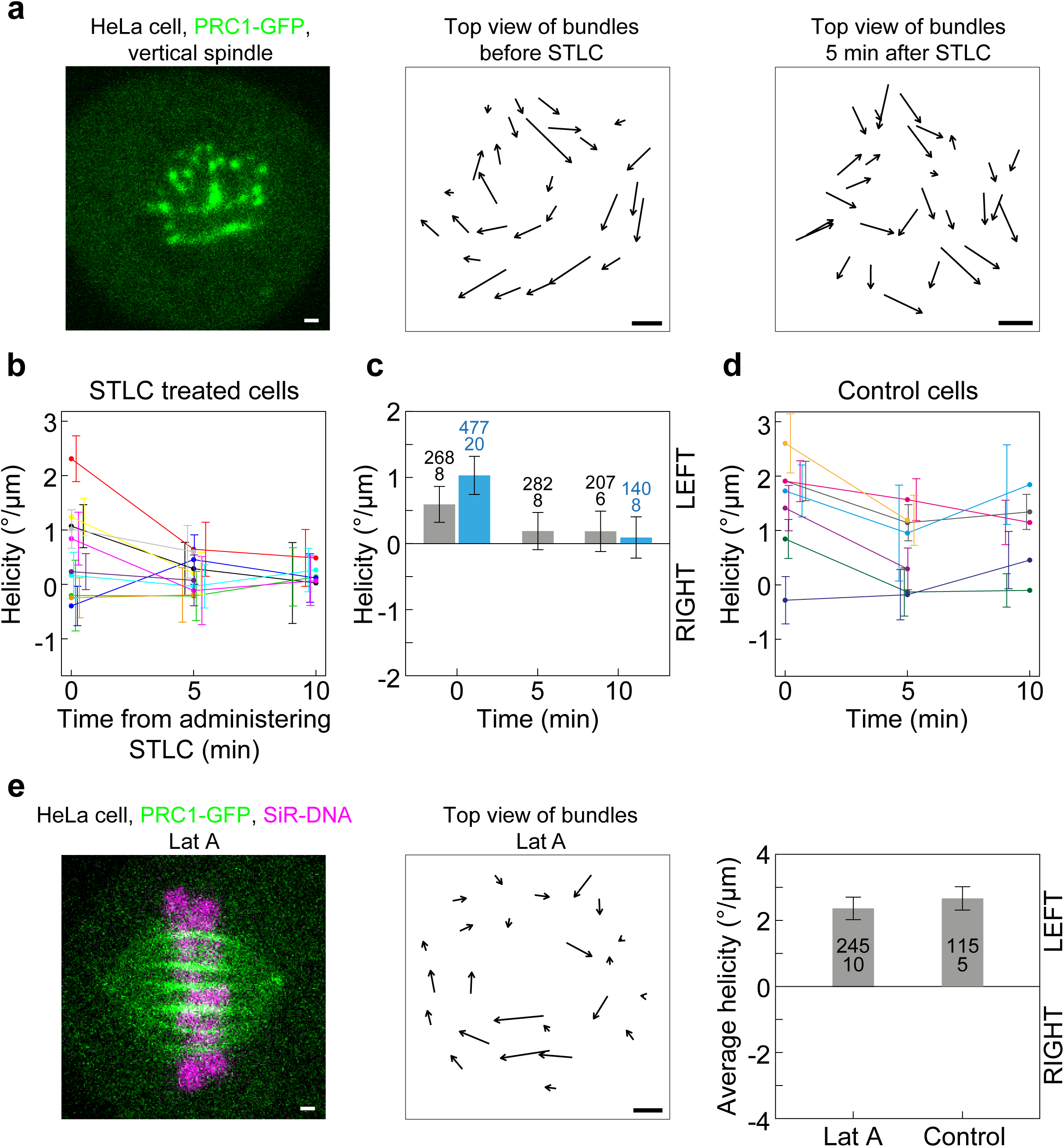
Perturbation of Eg5 reduces the spindle chirality, whereas perturbation of astral microtubules does not. **a,** Confocal image of the equatorial plane of the metaphase spindle in HeLa cell expressing PRC1-GFP (left); arrows connecting starting and ending points of PRC1-GFP bundles traced in the upwards direction from the same cell before (middle) and 5 minutes after treatment with STLC (right). Note that the bundles before treatment rotate clockwise, whereas after the treatment there is no obvious rotation. **b,** Spindle average helicity of individual HeLa cells expressing PRC1-GFP measured before and at 5 and 10 minutes after treatment with STLC, where number of cells was n=10, 10, 6, respectively. Each cell is represented by dots of different color which are connected with a line. **c,** Spindle helicity calculated over all bundles shown as the average change in angle with height. HeLa cell expressing PRC1-GFP (grey), U2OS cell expressing photoactivatable-GFP-tubulin, mCherry-tubulin and EGFP-CENP-A (blue) before treatment with STLC and at 5 and 10 minutes after treatment, respectively. Positive values denote left-handed helicity, negative values denote right-handed helicity. **d,** Spindle average helicity of individual HeLa cells expressing PRC1-GFP measured before and at 5 and 10 minutes after mock-treatment (DMSO). Each cell is represented by dots of different color which are connected with a line. **e,** Metaphase spindle of a HeLa cell expressing PRC1-GFP with chromosomes labelled with SiR-tubulin assembles normally in the presence of 2μM latrunculin A (LatA) (left); arrows connecting starting and ending points of PRC1-GFP bundles from the cell on the left traced in the upwards direction (middle); Spindle helicity calculated over all bundles shown as the average change in angle with height in HeLa cell expressing PRC1-GFP, treated with latrunculin A for 60 minutes and control cells (right). Positive values denote left-handed helicity, negative values denote right-handed helicity. The numbers in each bar represent the number of bundles (top) and cells (bottom), scale bars represent 1μm, error bars represent s.e.m.

Twist in the spindle may be also regulated by astral microtubules. To test the contribution of astral pulling forces at the cell cortex to the twist in the spindle, we treated the cells with latrunculin A, which depolymerizes cortical actin^19^. Consequently, it disrupts the cortical localization of LGN protein, which is a part of NuMA/LGN/Ga complex that anchors dynein on the cell cortex^20,21^. Cortical dynein positions the spindle with respect to the cell cortex by pulling on astral microtubules^22^. We found that microtubule bundles in latrunculin-treated Hela cells had a helicity of 2.4±0.4 °/μm (n=245 bundles from 10 cells, Fig. 2e), which is similar to the bundle helicity in untreated HeLa cells (2.7±0.4 °/μm, 115 bundles from 5 cells). In these experiments, we used horizontally oriented spindles to confirm the effect of latrunculin A on spindle position (Extended Data Fig. 2b and Methods). Our results indicate that pulling forces generated by astral microtubules at the cell cortex have a minor effect on spindle chirality.

To explain the observed shapes, we introduce a theoretical model for the balance of forces and torques acting on the microtubule bundles in the spindle, taking into account microtubule elastic properties. In our model, two spindle poles are represented as spheres of radius *d* with centers separated by vector **L** of length *L* = |**L**|, whereas microtubule bundles are represented as curved lines connecting these spheres (Fig. 3a). Microtubule bundles, denoted by index *i* = 1, …, *n*, extend between points located at the surface of the left and right sphere, where positions with respect to center of each sphere are given by vectors **d**_*i*_ and **d**_*i*_′, respectively. denotes number of microtubule bundles. Based on the observation that spindle shape is rather constant during metaphase, we introduce balance of forces and torques on the spindle pole. For the left pole, force balance reads

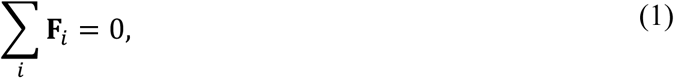

and the balance of torques reads

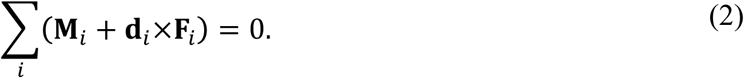

**Figure 3.**
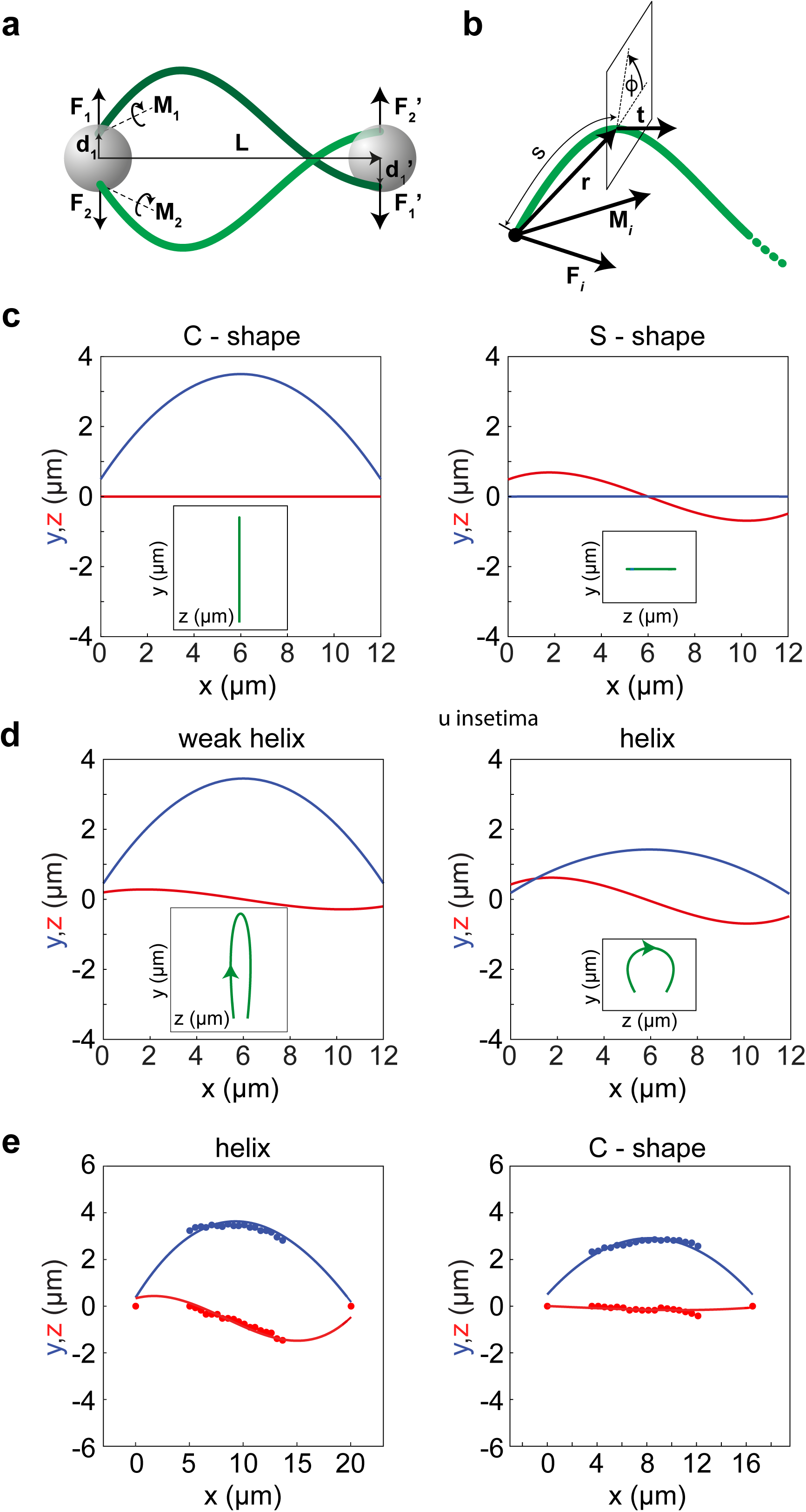
Theory of spindle shape. **a,** Scheme of the model. Microtubule bundles (green) extend between 2 spindle poles (grey spheres) at the distance **L**. Straight arrows denote forces **F**_1,2_ and positions at the spheres **d**_1,2_, whereas curved arrows denote torques **M**_1,2_. **b**, Scheme of a microtubule bundle. Arrows denote the contour length *s*, radial vector, **r**, the normalized tangent vector **t** and the torsion angle, *ϕ*. **c** and **d,** Predicted shapes of the microtubule bundles. Three different projections are shown: *xy* projection (blue), *xz* projection (red) and in insets *yz* projection (green). In **c** used parameters are **M**_1_ = (0, –150, 0) *pN*μ*m* and **M**_1_ = (0, 0, 150) *pN*μ*m* in left and right panel, respectively. **e**, Theoretical fits to the traces of microtubule bundles from live HeLa cells expressing PRC1-GFP. Examples of fits to helical shape (left) and C-shape (right) are shown. Used parameters are *M* = (–2.6, –74.6, 81.8) *pN*μ*m* for the helical shape and *M* = (–0.2, –0.7,64.0) *pN*μ*m* for the C-shape. In panels **c** and **d** parameter *L* = 12 μ*m*, whereas in **e**, *L* is taken from measurements. The other parameters are *d* = 0.5 μ*m* and *κ* = 900 *pN*μ*m*^2^.

Here, **F**_*i*_ and **M**_*i*_ denote forces and torques exerted by the left pole at the *i*–th microtubule bundle, respectively. Balances of forces and torques at the right pole are obtained by equations analogues to equations (1) and (2), where **F**_*i*_, **M**_*i*_, and **d**_*i*_ are replaced with **F**_*i*_′, **M**_*i*_′, and **d**_*i*_′, respectively. Here and throughout the text the prime sign corresponds to the right pole. We also introduce balance of forces and torques for the microtubule bundle

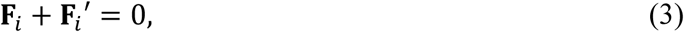

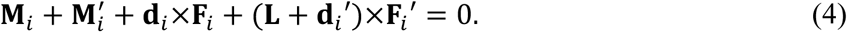

Forces and torques acting at the microtubule bundle change its shape, because microtubule bundles are elastic objects^13,23,24^. We describe a microtubule bundle as a single elastic rod of bending elasticity *κ* and torsional rigidity *τ*. The contour of elastic rod is described by its length, *s*, and radial vector, **r**(*s*). The normalized tangent vector is calculated by **t** = *d***r**/*ds*. The torsion angle, *ϕ*(*s*), describes orientation of the cross-section along the length of the rod (Fig. 3b). The curvature and the torsion of an elastic rod are described by the static Kirchoff equation^25^

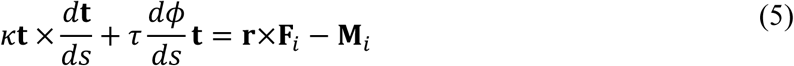

Our model provides a description for the force and torque balance of the entire spindle and makes a link between shape of microtubule bundles and forces and torques at the spindle poles.

The model describes a system consisting of *n* microtubule bundles, where torques and forces can vary between bundles, resulting in a system with a large number of degrees of freedom. To reduce the number of degrees of freedom, we consider a case with two microtubule bundles, *i* = 1, 2. Further, we use rotational symmetry of the spindle with respect to the major axis by imposing the symmetry for forces **F**_1∥_ = **F**_2∥_, **F**_1⊥_ = –**F**_2⊥_ and for torques **M**_1∥_ = **M**_2∥_, **M**_1⊥_ = –**M**_2⊥_. Here, index ∥ and ⊥ denotes components of vectors that are parallel and perpendicular to the vector **L**, respectively, obeying **F**_*i*_ = F_*i*∥_ + **F**_*i*⊥_ and **M**_*i*_ = **M**_*i*∥_ + **M**_*i*⊥_. In addition, we impose that the magnitude of torque is equal at both poles, |**M**_*i*_| = |**M**_*i*_′|, that the components of torque parallel to **L** are balanced, **M**_*i*∥_ = –**M**_*i*∥_′, and **d** · **M**_*i*⊥_ = **d**′ · **M**′_*i*⊥_ = 0. For simplicity, we also choose that vectors **d** and **d**′ are perpendicular to **L**. To solve the model, we choose a Cartesian coordinate system with the origin at the center of left spindle pole and *x*-axis parallel to **L**. In this coordinate system, radial vector has components **r** = (*x, y, z*) and torques have components **M**_*i*_ = (*M*_*ix*_, *M*_*iy*_, *M*_*iz*_). The orientation of the coordinate system is chosen such that *M*_*iy*_ = *M*_*iy*_′ and *M*_*iz*_ = – *M*_*iz*_′. Thus, our model has only two free parameters (see Methods).

In the small angle approximation, equation (5) has simple analytical solutions and the twisting moment corresponds to the component of torque parallel to *x*-axis. If torque has a bending moment only, *M*_1*x*_ = 0, there are two solutions which are both planar, the symmetric C-shape and the anti-symmetric S-shape (Fig. 3c). However, by adding a twisting moment, shapes become three-dimensional with a non-vanishing helicity in either case (Fig. 3d). Thus, our theory predicts that a twisting moment is required for a microtubule bundle to have a helical shape.

To compare the results of our model with the experimentally observed shapes of microtubule bundles, we fit our analytical solutions to the traces of microtubule bundles from live HeLa cells expressing PRC1-GFP (Fig. 3e). Our theory reproduces three-dimensional helical shapes, as well as symmetric C-shapes (Fig. 3e; Extended Data Fig. 3 shows additional examples). In the case of the helical shape shown in Fig. 3e the twisting moment was *M*_1*x*_ = –2.6 *pN*μ*m*, whereas in the case of the C-shapes it was 10 times smaller. The bending moment was similar in both cases. For 48 bundles the twisting moment was *M*_1*x*_ = –0.5 ± 0.2 *pN*μ*m* and the bending moment was |**M**_1⊥_| = 69 ± 4 *pN*μ*m* (see Methods). Our results show that a twisting moment is required to reproduce the experimentally observed helical shapes, whereas a bending moment is required for all curved shapes.

In summary, we found that the mitotic spindle is a chiral object. Chirality is an intriguing property of the biological world, present at all scales ranging from molecules to whole organisms. We find that spindle chirality cannot be explained by forces but rather by torques. Based on our experiments and theory, we conclude that torques generated by motor proteins, in addition to forces, exist in the spindle and determine the spatial organization of the microtubule bundles.

## Methods

### Cell culture and sample preparation

HeLa-Kyoto BAC lines stably expressing PRC1-GFP were courtesy of Ina Poser and Tony Hyman (MPI-CBG, Dresden). HeLa cells stably expressing EGFP-CENP-A and EGFP-centrin1 were a courtesy from Emanuele Roscioli and Andrew McAinsh (WMS - Cell and Development Biology - University of Warwick). Human U2OS cells, permanently transfected with CENP-A-GFP, mCherry-α-tubulin and photoactivatable (PA)-GFP-tubulin, were courtesy from Marin Barišić and Helder Maiato (Institute for Molecular Cell Biology, University of Porto, Portugal). Cells were grown in DMEM with Ultraglutamine (Lonza, Basel, Switzerland), supplemented with 10% FBS, penicillin, streptomycin and geneticin (Santa Cruz Biotechnology, Inc., Dallas, USA). The cells were kept at 37°C and 5% CO_2_ in a Galaxy 170 R CO_2_ humidified incubator (Eppendorf, Hamburg, Germany).

To visualize kinetochores and identify the metaphase, HeLa celles expressing PRC1-GFP cells were transfected by electroporation using Nucleofector Kit R (Lonza, Basel, Switzerland) with the Nucleofector 2b Device (Lonza, Basel, Switzerland), using the high-viability O-005 program. Transfection protocol provided by the manufacturer was followed. 25-35 h before imaging, 1x10^6^ cells were transfected with 2.5 μg of mRFP-CENP-B plasmid DNA (pMX234) provided by Linda Wordeman (University of Washington). To visualize chromosomes and determine the metaphase state, 1 hour prior to imaging SIR-DNA (Spirochrome AG, Stein am Rhein, Switzerland) was added to the dish with live HeLa cells at 100nM final concentration.

To prepare samples for microscopy, HeLa and U2OS cells were seeded and cultured in 1.5 mL DMEM medium with supplements at 37°C and 5% CO_2_ on uncoated 35-mm glass coverslip dishes, No 1.5 coverglass (MatTek Corporation, Ashland, MA, USA). Before live-cell imaging, the medium was replaced with Leibovitz’s L-15 CO_2_-independent medium supplemented with fetal bovine serum (FBS, Life Technologies, Carlsbad, CA, USA). For experiments with the fixed samples, cells were fixed in ice-cold methanol for 3 min and washed three times with phosphate buffered saline (PBS, Merck, Darmstadt, Germany).

### Drug treatments

The stock solution of S-trityl-L-cysteine (STLC) and latrunculin A were prepared in DMSO to a final concentration of 1mM. Both drugs and solvent were obtained from Sigma-Aldrich. The working solution was prepared in DMEM medium at 2x final concentration. At the time of treatment, the working solution was added to cells at 1:1 volume ratio. STLC treated samples were acquired as follows: images of a cell with vertical spindle were acquired, then the drug was added at a final concentration of 50 μM and the same spindle was imaged after 5 and 10 minutes. Appearance of monopolar spindles confirmed the effect of STLC. For latrunculin A treatment experiment, the PRC1-GFP HeLa cells were treated with 2 μM latrunculin A for one hour prior to imaging, which was done between one and two hours post treatment. The effect of latrunculin A was confirmed by significant cell blebbing immediately after drug addition, decrease of cell diameter and spindle mis-positioning (Extended Data Fig. 2b). Here 21 out of 30 latrunculin-treated cells had spindles close to the cell cortex, which is rare in untreated cells). For mock-treated experiments, cells were treated with the concentration of DMSO that was used for preparation of drug treatments.

### STED microscopy

STED images of HeLa and U2OS cells were recorded at the Core Facility Bioimaging at the Biomedical Center, LMU Munich. STED resolution images were taken of Sir-Tubulin signal, whereas GFP signal of kinetochores and centrin1 was taken in confocal resolution. Gated STED images were acquired with a Leica TCS SP8 STED 3X microscope with pulsed White Light Laser excitation at 652 nm and pulsed depletion with a 775 nm laser (Leica, Wetzlar, Germany). Objective used was HC PL APO CS2 93x/1.30 GLYC with motorized correction collar. Scanning was done bidirectionally at 30-50 Hz, a pinhole setting of 0.93 AU (at 580 nm) and the pixel size was set to 20 x 20 nm. The signals were detected with Hybrid detectors with the following spectral settings: Sir-Tubulin (excitation 652; emission: 662 - 715 nm; counting mode, gating: 0.35 - 6 ns), GFP (excitation 488; emission 498-550; counting mode, no gating).

### Confocal microscopy image acquisition

Fixed HeLa cells expressing PRC1-GFP were imaged by using a Leica TCS SP8 X laser scanning confocal microscope with a HC PL APO 63x/1.4 oil immersion objective (Leica, Wetzlar, Germany) heated with an objective integrated heater system (Okolab, Burlingame, CA, USA). Excitation and emission lights were separated with Acousto-Optical Beam Splitter (AOBS, Leica, Wetzlar, Germany). For excitation, a 488-nm line of a visible gas Argon laser and a gated STED supercontinuum visible white light laser at 575 nm were used for GFP and mRFP, respectively. GFP and mRFP emissions were detected with HyD (hybrid) detectors in ranges of 498–558 and 585–665 nm, respectively. Pinhole diameter was set to 0.8 μm. Images were acquired at 25-35 focal planes with 0.5 μm spacing and 400 Hz unidirectional xyz scan mode. The system was controlled with the Leica Application Suite X software (LASX, 1.8.1.13759, Leica, Wetzlar, Germany).

Live HeLa and U2OS cells were imaged using Bruker Opterra Multipoint Scanning Confocal Microscope (Bruker Nano Surfaces, Middleton, WI, USA). The system was mounted on a Nikon Ti-E inverted microscope equipped with a Nikon CFI Plan Apo VC 100x/1.4 numerical aperture oil objective (Nikon, Tokyo, Japan). During imaging, cells were maintained at 37°C in Okolab Cage Incubator (Okolab, Pozzuoli, NA, Italy). 60 μm pinhole aperture was used and the xy-pixel size was set to 0.83 μm by placing a 2x relay lens in front of the camera. For excitation of GFP and mCherry fluorescence, a 488 and a 561 nm diode laser line were used, respectively. The excitation light was separated from the emitted fluorescence by using Opterra Dichroic and Barrier Filter Set 405/488/561/640. Images were captured with an Evolve 512 Delta EMCCD Camera (Photometrics, Tucson, AZ, USA) with no binning performed. To cover the whole metaphase vertical spindle, Z-stacks were acquired at 30-60 focal planes separated by 0.5 μm spacing with unidirectional xyz scan mode and without frame averaging. In latrunculin A treated cells horizontal spindles were imaged, with Z-stacks acquired at 48 focal planes separated by 0.5 μm spacing with unidirectional xyz scan mode and with four frame averages. The system was controlled with the Prairie View Imaging Software (Bruker Nano Surfaces, Middleton, WI, USA).

### Image analysis

Microscopy images were analysed in Fiji software^26^. Tracking of bundles was done using the available Multi-point tool. Obtained data was further analysed and plotted in R programming language.

Transformation of vertical spindles from horizontal spindle images was also done using code written in R. Since images obtained by using confocal microscopy have a gap between them, during transformation we had to fill this gap by copying the same pixel a required number of times depending on the initial pixel size.

Also, the gap size doesn’t have to be a multiple of the pixel size. To take this into consideration, pixel width was changed accordingly. For instance, if we had a 500 nm gap between horizontal spindle images and 80 nm pixels, the best we could do is copy 6 pixels in the gap. This would leave us with unaccounted 20 nm so we introduced a correction by saying the width of each pixel is 500/6 = 83.3 nm. To ensure that spindles are maximally vertical after the transformation, horizontal spindles were chosen for transformation if both poles could be seen simultaneously in one image. For the same reason, before transformation, the images were rotated so that the spindle long axis was approximately parallel to the x axis. Analogously, vertical spindle images were transformed into a horizontal spindle.

### Forces and torques for the system of 2 bundles and with the imposed symmetries

From equation (1) and the symmetry **F**_1∥_ = **F**_2∥_, we obtain that, *F*_1*x*_ = *F*_2*x*_ = 0. From **F**_1⊥_ = –**F**_2⊥_, the other two components obey *F*_1y_ = –*F*_2*y*_, *F*_1*z*_ = –*F*_2*z*_. For coordinate system with orientation in which *M*_*iy*_ = *M*_*iy*_′ and *M*_*iz*_ = –*M*_*iz*_′ the *z*–component of equation (4) reads *LF*_1*y*_ = 0 and *y*–component of the force vanisfes, *F*_1*y*_ = 0. By using the *x*–component of equation (2), *M*_1*x*_ + *d*_*y*_*F*_1*z*_ = 0, together with the *y*–component of equation (4), 2*M*_1*y*_ – *LF*_1*z*_ = 0, we obtain *d_y_* = –*M*_1*x*_*L*/2*M*_1*y*_. By using the *x*–component of equation (2) for the right centrosome, we obtain *d_y_* = *d_y_*′. Because we imposed the symmetry **d** · **M**_1⊥_ = **d**′ · **M**_1⊥_′ = 0, and **d** is perpendicular to **L**, components of vectors **d** and **d**′ obey *d_z_* = –*d_y_M*_1*y*_/*M*_1*z*_ and *d_z_* = –*d*′_*z*_. By using 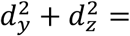 = *d*^2^, we obtain the relation between the parameters *M*_1*x*_, *M*_1*y*_ and *M*_1*z*_:

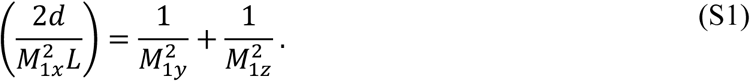

### Analytical solutions of equation (5)

To solve equation (5) we use a Cartesian coordinate system in which this equation is given by system of three nonlinear differential equations. In the small angle approximation, where *ds* ≈ *dx*, these equations simplify and become linear:

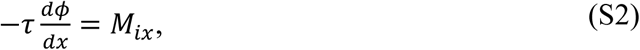

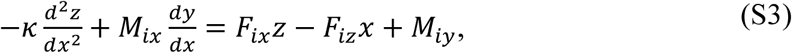

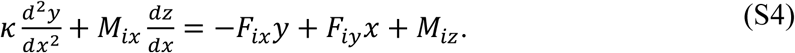

Analytical solutions of equations (S2)-(S3) reads:

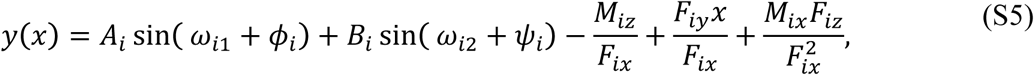

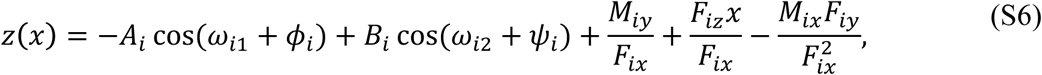

where 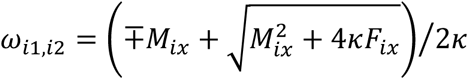.

Note that in the case of two bundles, with imposed symmetries, these solutions simplify because *F*_1*x*_ = *F*_1*y*_ = 0. Integration constants *A*_1_, *B*_1_, *ϕ*_1_, *ψ*_1_ are obtained from the boundary conditions *y*(0) = *y*(*L*) = *d_y_*, *z*(0) = –*z*(*L*) = *d_z_*

### Comparison of the model to experimentally observed shapes

We have compared the theoretically obtained shapes, given by equations (S5) and (S6), to the tracking data of live HeLa cells expressing PRC1. The parameters of the fit are *M*_*x*_ and *M*_*y*_, and the orientation of the coordinate system of the tracked shape. Used parameters are *d* = 0.5 μ*m*, and *κ* = 900 *pN*μ*m*^2^. Parameter *L* is obtained from the experimentally measured distance between the poles. Torque *M*_*z*_ and forces *F*_*y*_ and *F*_*z*_ are calculated after all other parameters are known. We fitted 60 traced bundles and for 80 % of all the shapes discrepancy between fitted curves and experimental data was smaller than:

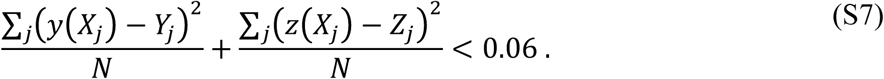

Here, *X*_*j*_, *Y*_*j*_, *Z*_*j*_ are measured coordinates of the imaging plane *j*, and *N* denotes number of data points used for fitting a single bundle. We used shapes with maximal distance from the major axis larger than 1 μ*m*.

## Acknowledgments

We thank Andrew McAinsh, Emanuele Roscioli, Ina Poser, Tony Hyman, Marin Barišić and Helder Maiato for cell lines; Igor Weber, Maja Marinović, Vedrana Filić Mileta and the rest of the Weber lab for help with the confocal microscope. We thank Steffen Dietzel, Anna H. Klemm and the Core Facility Bioimaging at the Biomedical Center – LMU, Munich, Germany for help with STED microscopy. We thank Ivana Šarić for the drawings. We thank all the members of Tolić and Pavin labs for helpful discussions. J.S. and M.N. acknowledge support from the European Social Fund (INTERBIO); M.N. acknowledges support from Unity through Knowledge Fund (UKF); N.P. acknowledges support from QuantiXLie Center of Excellence; I.M.T. acknowledges support from European Research Council (ERC Consolidator Grant, GA Number 647077).

## Author contribution

M.N. developed the theoretical model. J.S. and B.P. performed confocal microscopy experiments. Z.B. analyzed the experimental data with help from B.K. J.S. and B.K. carried out STED imaging, with A.T. providing expertise on STED microscopy. N.P and I.M.T. conceived the project and supervised theory and experiments, respectively. M.N., J.S., I.M.T., and N.P wrote the paper with input from all authors.

## Extended Data

**Extended Data Figure 1**

**a**, STED image of a metaphase spindle in a U2OS cell expressing photoactivatable-GFP-tubulin, mCherry-tubulin and EGFP-CENP-A. Different shapes of microtubule bundles (white tracks) extend almost through the whole spindle (left).

**b** and **c**, arrows connecting starting and ending points of PRC1-GFP bundles fixed HeLa cells with vertical, **b**, and horizontal, **c**, spindles traced in the upwards direction. Note that the bundles rotate clockwise, revealing left-handed chirality of the spindle.

**d,** Bars show spindle helicity as the average change in angle with height, calculated over cells where helicity of an individual cell is calculated by considering all bundles from the respective cell. Positive values denote left-handed helicity, negative values denote right-handed helicity. Data are shown for HeLa cells expressing PRC1-GFP, and U2OS cells expressing CENP-AGFP, mCherry-α-tubulin and photoactivatable (PA)-GFP-tubulin. The numbers in each bar represent the number of bundles (top) and cells (bottom), error bars represent s.e.m. Scale bars in panels **a-c** represent 1μm.

**Extended Data Figure 2**

**a,** Spindle helicity shown as the average change in angle with height, averaged over cells. HeLa cell expressing PRC1-GFP (grey), U2OS cell expressing photoactivatable-GFP-tubulin, mCherry-tubulin and EGFP-CENP-A (blue) before treatment with STLC and at 5 and 10 minutes after treatment, respectively. Positive values denote left-handed helicity, negative values denote right-handed helicity.

**b,** Metaphase spindle of a HeLa cell expressing PRC1-GFP (green) with chromosomes labelled with SiR-DNA. Addition of 2μM Latrunculin A causes blebbing of the cell membrane (left) and mispositioning of the spindle (right). The numbers in each bar represent the number of bundles (top) and cells (bottom), scale bars, 1μm, error bars represent s.e.m.

**Extended Data Figure 3**

Theoretical fits to the traces of microtubule bundles from live HeLa cells expressing PRC1-GFP. Examples of fits to helical shape (left) and C-shape (right) are shown. Fitted parameters are shown in figures. Parameter *L* is taken from measurements, *d* = 0.5 μ*m* and *κ* = 900 *pN*μ*m*^2^.

**Supplement Video 1**

PRC1-labeled microtubule bundles rotate around the spindle axis. Scheme of imaging of a vertically oriented spindle (left); images of a vertically oriented spindle in a HeLa cell expressing PRC1-GFP (right) corresponding to the planes shown on the scheme. Note that the bundles rotate clockwise as the imaging plane moves upwards, revealing the left-handed chirality of the spindle.

**Supplement Video 2**

Three-dimensional projections with the coordinate system represented as a cuboidal box. Dots represent traced bundles from HeLa cells expressing PRC1-GFP. Colors represent different bundles.

